# Development of a high throughput system to screen compounds that revert the activated hepatic stellate cells to a quiescent-like state

**DOI:** 10.1101/2023.11.01.564270

**Authors:** Yasuhiro Nakano, Tohru Itoh, Minoru Tanaka, Atsushi Miyajima, Taketomo Kido

## Abstract

Chronic liver injury induces fibrosis that often proceeds to cirrhosis and hepatocellular carcinoma, indicating that prevention and/or resolution of fibrosis is a promising therapeutic target. Hepatic stellate cells (HSCs) are the major driver of fibrosis by expressing extracellular matrices (ECM). HSCs, in the normal liver, are quiescent and activated by liver injury to become myofibroblasts that proliferate and produce ECM. It has been shown that activated HSCs (aHSCs) become a “quiescent-like” state by removal of liver insults. Therefore, deactivation agents can be a therapeutic drug for advanced liver fibrosis. Using aHSCs prepared from human induced pluripotent stem cells, we found that aHSCs were reverted to a quiescent-like state by a combination of chemical compounds that either inhibit or activate a signaling pathway, Lanifibranor, SB431542, Dorsomorphin, retinoic acid, palmitic acid and Y27632, *in vitro*. Based on these results, we established a high throughput system to screen agents that induce deactivation and demonstrate that a single chemical compound can induce deactivation.

## Introduction

Fibrosis is characterized by excessive accumulation of extracellular matrix (ECM) in chronically injured organs and is an essential process for tissue repair by encapsulating the damaged area^1^. In the liver, chronic inflammation induced by a variety of etiologies, including hepatitis virus infection, alcohol abuse, and nonalcoholic steatohepatitis (NASH), results in fibrosis that often advances to cirrhosis and carcinogenesis^2^. Parenchymal hepatocytes are the major initial target of those liver injury and damaged hepatocytes release factors that activate hepatic stellate cells (HSCs) directly and indirectly via nonparenchymal cells such as macrophages^3,4^. HSCs in the normal liver are quiescent (qHSCs) and exhibit dendritic morphology and activated HSCs (aHSCs) play a central role for fibrosis by producing ECM^3,4^. TGF-β and PDGF are the two major profibrotic cytokines and activate HSCs in injured liver. Activated HSCs (aHSCs) are myofibroblastic cells that proliferate and produce ECM^3,4^.

While a large effort has been made to develop drugs for hepatitis such as NASH, they are mostly targeting hepatocytes but not directly target fibrosis^5^. Although some drugs that target hepatocytes also ameliorate fibrosis, their anti-fibrotic effect is limited. Cirrhosis is an advanced stage of fibrosis, at which no effective therapeutic option is available other than liver transplantation^6^. Liver fibrosis was historically considered to be a passive and irreversible process due to the substitution of damaged hepatic parenchyma with a collagen-rich tissue^2^. However, recent studies have shown histological improvement of liver fibrosis after removal of causative agents^4^. For example, cessation of carbon tetrachloride administration was shown to revert the aHSCs to a quiescent state, known as deactivation ^7,8^. Tcf21 has been identified as a deactivation factor of aHSCs. Adeno-associated virus-mediated expression of Tcf21 in aHSCs not only suppressed collagen expression but also restored cells, at least partly, to a quiescent-like phenotype, *in vitro* and *in vivo*. Those qHSC-like cells from aHSCs contribute to the improvement of hepatic architecture and function^9^. These results suggest that agents with a potential to revert aHSCs to qHSC-like cells should provide an excellent therapeutic option to advanced fibrosis.

A major problem to develop anti-fibrotic agents is the lack of appropriate HSC for drug discovery; HSC cell lines have lost characteristics of HSCs and it is practically impossible to prepare fresh normal HSC from the body for drug screening. We have previously established a protocol to induce differentiation of quiescent-like HSCs from human induced pluripotent stem cells (hiPSCs) that can be activated *in vitro* and demonstrated that those qHSCs can be used to screen anti-fibrotic drugs^10^. In this study, we show that the iPSC-derived aHSCs can be reverted to a pre-activation stage *in vitro* by a combination of chemical compounds that either inhibit or activate signaling pathways. We then establish a high throughput system (HTS) to screen deactivation drugs. Finally, we show that a single compound screened by the HTS can convert aHSCs to a quiescent-like stage.

## Results

### Deactivation of activated HSCs *in vitro*

As we reported previously, qHSCs and aHSCs can be generated from hiPSCs (Figure 1A). HSCs differentiated from hiPSCs using our protocol exhibited phenotype, similar to qHSC; expression of a number of qHSC marker genes including *NGFR*, *LRAT* and *NES*, and accumulation of vitamin A, but the lack of the expression of activation marker genes such as *ACTA2* (encoding α-SMA) and *COL1A1*^10^. Those qHSC-like cells were activated spontaneously in culture and expressed αSMA and collagens and diminished expression of quiescent marker genes. In addition, morphology of HSC significantly changed from dendritic qHSC-like cells to myofibroblastic aHSCs. With these changes, filamentous actin (F-actin) accumulation was observed in the cytoplasm visualized by phalloidin (Figure 1B), suggesting that HSC activation can be evaluated with F-actin.

**Figure 1.**
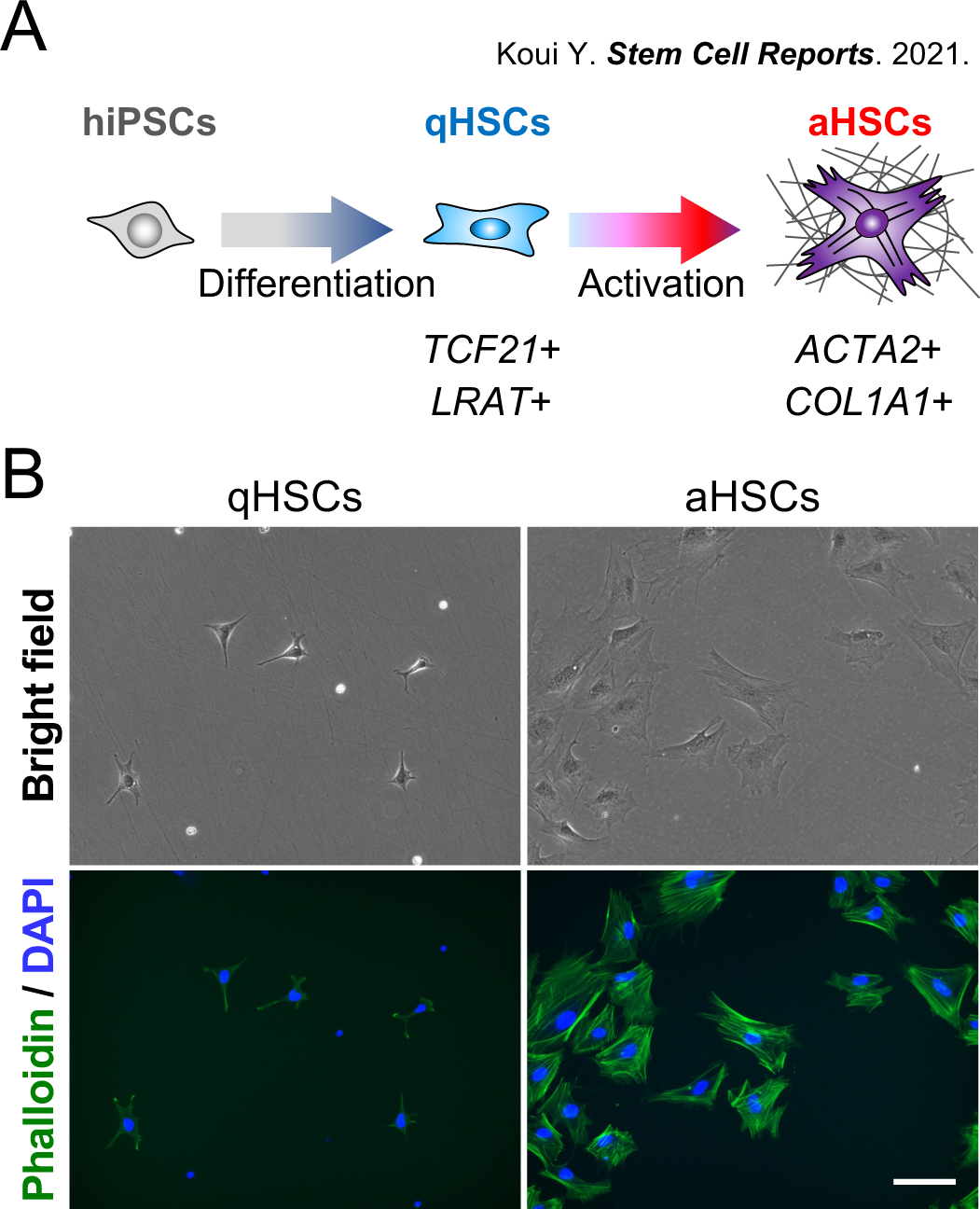
Generation of qHSCs and aHSCs from hiPSCs. (A) Preparation of qHSCs and sHSCs from hiPSCs. (B) Phase-contrast images of qHSCs and aHSCs (upper). Fluorescence images of qHSCs and aHSCs incubated with phalloidin (lower). Nuclei were stained with DAPI. Scale bar, 100 μm.

In order to establish a system to screen compounds that deactivate aHSCs, we examined whether hiPSC-derived aHSCs can be deactivated. Previous studies reported several potential chemical compounds that reverted aHSCs to a stage, similar to, but distinct from, the qHSCs. Using hiPSC-derived aHSCs, we evaluated chemical compounds that were reported to act as an inhibitor of activation HSCs or a possible deactivation factor; a pan-PPAR agonist Lanifibranor (L), retinoic acid (R), palmitic acid (P), TGF-β inhibitor SB431542 (S), AMPK inhibitor Dorsomorphin (D), and ROCK inhibitor Y27632 (Y). None of those compounds alone failed to induce deactivation. However, combinations of YLRP, SDRP, SDYL, SDYRP, or SDYLRP significantly suppressed the expression of *ACTA2* and *COL1A1*, and SDYL or SDYLRP induced the expression of *TCF21*, an important transcription factor that was shown to be involved in deactivation (Figure 2A). SDYLRP most significantly changed the gene expression. In order to evaluate the changes of gene expression more comprehensively, we performed RNA-sequence (RNA-seq) analysis of qHSCs (day 0), aHSCs (day 7, day 10) and deactivated HSCs (daHSCs) (day 10) by the addition of SDYLRP. RNA-seq analysis showed that *LHX2*, *LRAT* and *TCF21* were highly expressed in daHSCs, whereas *ACTA2* and *COL1A1* were down-regulated compared with aHSCs (Figure S1A). Pearson correlation coefficient was calculated to investigate the correlation in gene expression profile between the samples (Figure 2B). The results showed that daHSCs (day 10) and aHSCs (day 10) were highly correlated with qHSCs (day 0) and aHSCs (day 7), respectively (Figure 2B). This suggested that aHSCs at day 7 may have partially reverted to the quiescent phase after 3 days of incubation with SDYLRP. In addition to gene expression analysis, morphological changes were examined. The dendritic shape of qHSCs was changed to flat myofibroblastic morphology in aHSCs upon culture, which was returned to the dendritic shape by the addition of SDYLRP but not with DMSO (Figure 2C and Figure S1B). Phalloidin staining visualized that actin stress fibers induced by the activation was lost upon incubation with SDYLRP (Figure 2D). Based on these results, we considered that hiPSC-derived HSCs were deactivated by SDYLRP and were useful cells to screen deactivation agents, and F-actin staining was a useful indicator to evaluate the activation status of HSCs.

**Figure 2.**
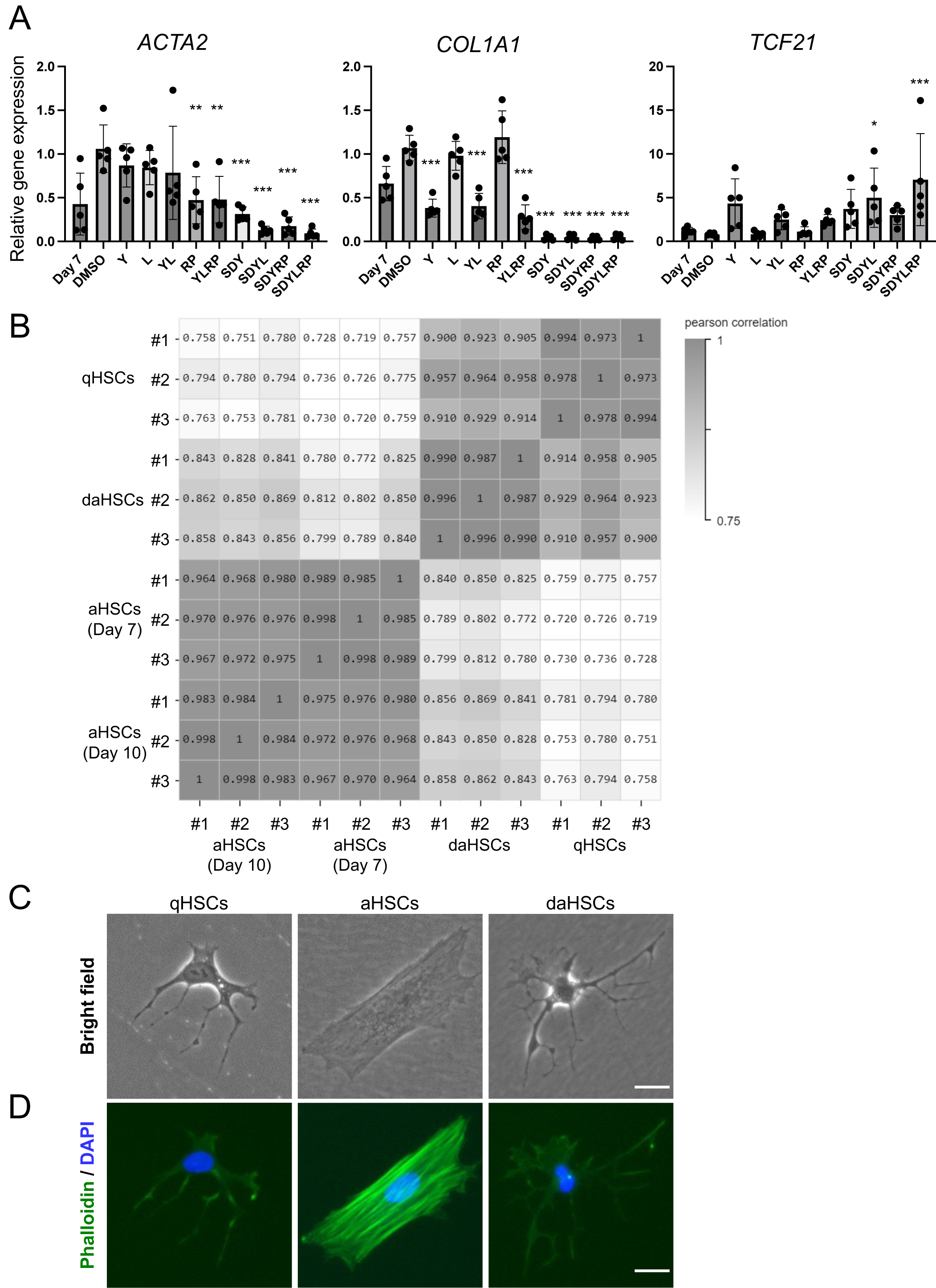
Deactivation of activated HSCs in vitro. (A) Expression levels of aHSC markers (*ACTA2* and *COL1A1*) and deactivation marker (*TCF21*). The results are shown as the mean ± SD of five independent experiments (each experiment contains three technical replicates). Y: Y27632, L: Lanifibranor, R: Retinoic acid, P: Palmitic acid, S: SB431542, D: Dorsomorphin. The expression level of 10-day cultured aHSCs without DMSO was set to 1. *P<0.05, **P<0.01, ***P<0.001. (B) Pearson correlation coefficient of gene expression profiles between qHSCs, aHSCs (day 7), aHSCs (day 10) and daHSCs (day 10) (C) Phase-contrast images of qHSCs, aHSCs and daHSCs. Scale bar, 25 μm. (D) Fluorescence images of qHSCs, aHSCs (day 10) and daHSCs incubated with phalloidin. Nuclei were stained with DAPI. Scale bar, 25 μm.

### Development of a high throughput system to screen deactivation agents

Since high throughput screening (HTS) requires a large number of aHSCs, we set up a large-scale preparation of aHSCs from hiPSCs and confirmed aHSCs maintained their activated phenotype after cryopreservation. Thawed cells were seeded in 384-well plates and incubated for 3 days with test compounds. We validated the assay system by using SDYLRP as a positive control and stained the cells with silicon-rhodamine (SiR)-Actin that was used to visualize F-actin in living cells^11^ (Figure 3A). The fluorescence intensity and the stained area were measured (Figure 3B). To quantitatively assess the activation state, the SiR-Actin fluorescence intensity per total cellular area (SiR-Actin intensity / SiR-Actin (+) area) was calculated as “Actin score” for each well, and plotted in Figure 3C. Quantification of the fluorescence intensity by an image cytometer showed that the wells with DMSO met the HTS validation criteria, i.e. the CV value was less than 10% and the Z’-factor was higher than 0.5^12,13^, indicating that the screening system is stable and has minimum sample-to-sample errors (Figure 3C). The fluorescence intensity was markedly declined by SDYLRP (Figure 3B), and Actin score was clearly distinguished from the well with DMSO (Figure 3C). Thus, we established a HTS system to screen deactivation agents.

**Figure 3.**
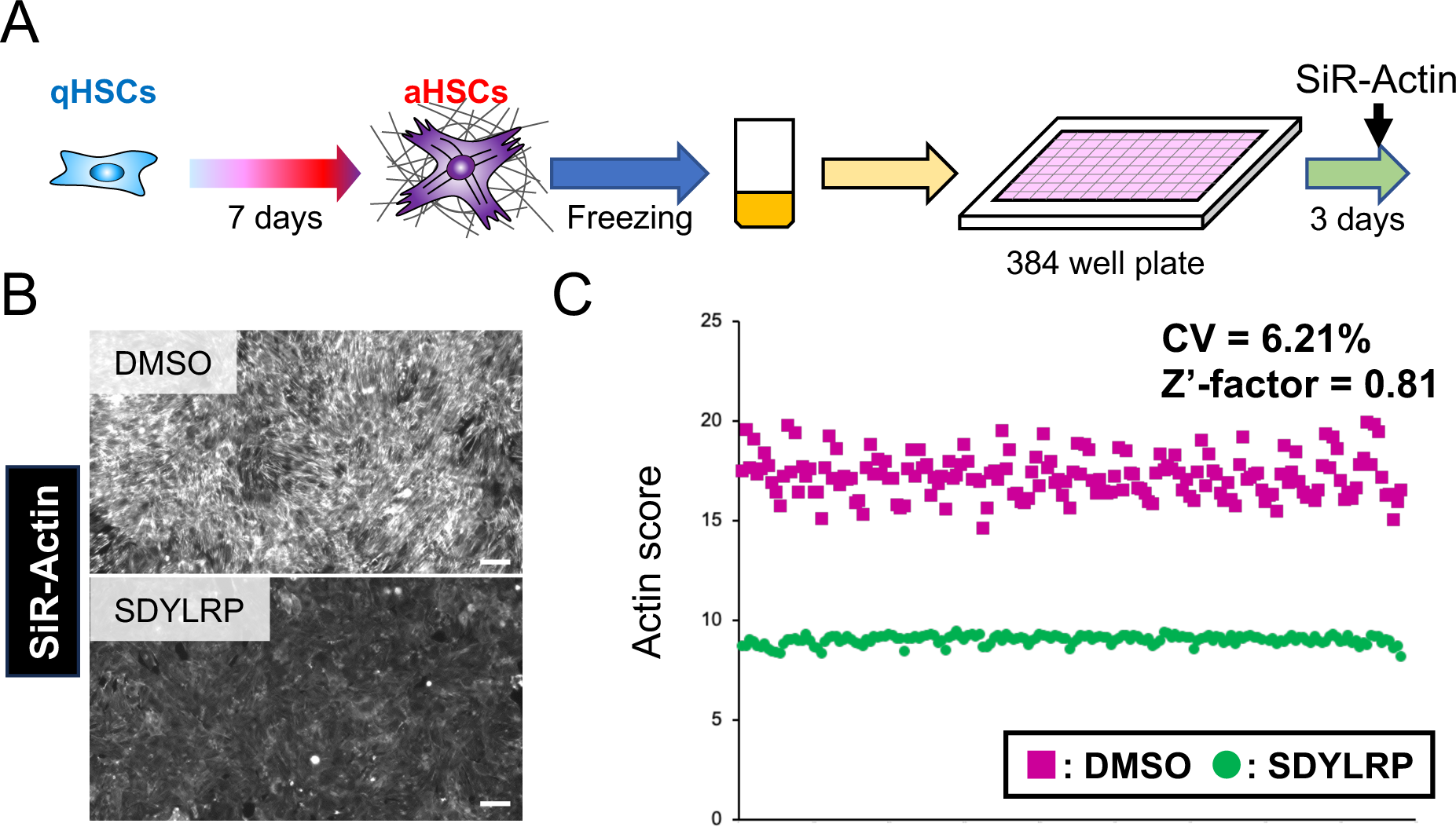
Development of a high throughput system to screen deactivation agents. (A) Schematic image of a high throughput system to screen deactivation agents. (B) Fluorescence images of F-actin accumulation using SiR-actin staining of HSCs incubated with DMSO or SDYLRP. Scale bar, 50 μm. (C) Dot plots of Actin score per well of 384-well plates incubated with DMSO or SDYLRP.

We applied this system to find deactivation chemicals, by fluorescent intensity, in the validated chemical library. Actin score of each well was quantified on a plate-by-plate basis and we selected the wells that exhibited fluorescence intensities below -2 × standard deviation (SD) on average in each plate. In the validated chemical library consisting of 3,909 compounds, we selected 171 compounds as first candidates that were effective at 2 μM. As a secondary screening, to select compounds with high deactivation activity, we tested the compounds at the concentration of 400 nM, and selected 25 compounds in the same plate (Figure 4A, 4B and Figure S2). We then performed a tertiary screening by quantitative PCR. The expression levels of the activation marker genes *ACTA2* and *COL1A1*, the deactivation gene *TCF21*, and the qHSC marker genes *LRAT* and *LHX2* are examined at a concentration of 125 nM, 500 nM and 2 μM (Figure S3) and are shown as a heatmap in Figure 4C. The expression level was set as 1 for each gene when DMSO was added as a negative control. Almost all compounds suppressed expression of *ACTA2* and *COL1A1* and increased *TCF21*, *LRAT* and *LHX2* expression at a concentration of 2 μM. This indicates that the HTS used in the primary and secondary screening is useful for finding compounds that induce deactivation. Some compounds showed a pattern of deactivation not only at 2 μM, but also at both 500 nM and 125 nM.

**Figure 4.**
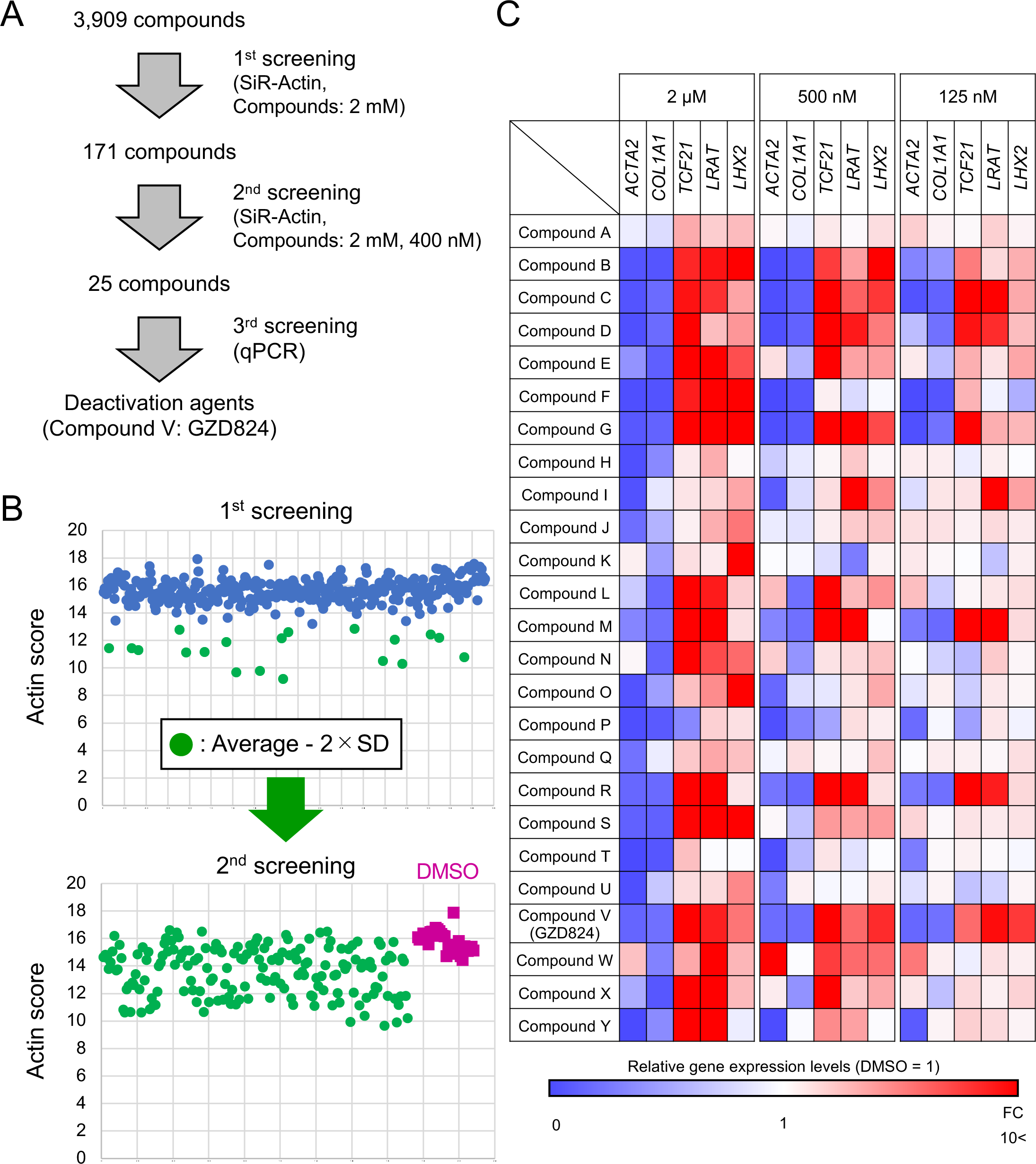
Screening of deactivation chemicals from the validated chemical library. (A) Summary of the results of drug screening. (B) Dot plots of Actin score per well of 384-well plates incubated with compounds of the chemical library. (C) Heat map of the relative expression levels of *ACTA2*, *COL1A1*, *TCF21*, *LRAT* and *LHX2* in hiPSC-derived aHSCs incubated with compound A to Y at the concentration of 125 nM, 500 nM and 2 μM. The values of samples treated with DMSO were set to 1.

### Deactivation of aHSC by a single selected compound

Among those 25 selected compounds, we have chosen Compound V, which corresponds to GZD824 (also known as Olverembatinib), a pan-Bcr-Abl inhibitor, to further evaluate the deactivation activity. GZD824 inhibited the fibrosis markers *ACTA2* and *COL1A1* in a concentration-dependent manner, while inducing the expression of the quiescent markers *LHX2* and *LRAT* as well as liver regeneration factors, *MDK* and *PTN*, in hiPSC-derived aHSCs (Figure 5A, 5B, 5C)^3,14^. Importantly, *TCF21*, a key transcription factor for aHSC deactivation, was dramatically up-regulated (Figure 5D), and along with this change in gene expression, cell morphology changed from aHSCs to qHSC-like cells with diminished F-actin accumulation (Figure 5E). This strongly suggested that GZD824 induced the deactivation of aHSCs, and daHSCs could improve liver function as well as fibrosis.

**Figure 5.**
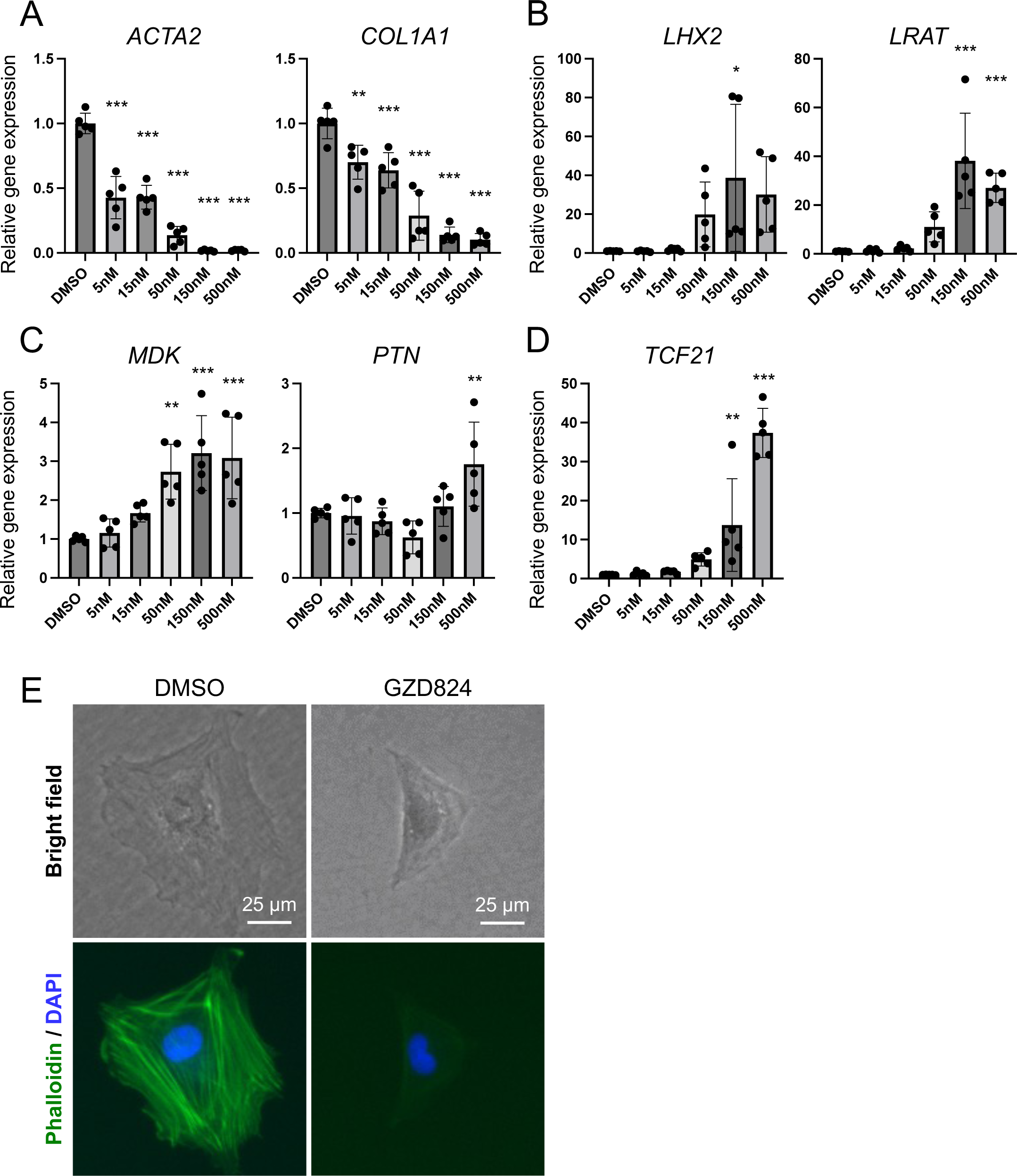
Deactivation of aHSC by GZD824. (A-D) Expression levels of (A) aHSC markers, (B) qHSC markers, (C) liver regeneration markers and (D) deactivation factor. The results are shown as the mean ± SD of independent experiments (each experiment contains three technical replicates). *P<0.05, **P<0.01, ***P<0.001. (E) Phase-contrast images (upper) and fluorescent phalloidin-stained images (lower) of aHSCs incubated with DMSO or GZD824. Nuclei were stained with DAPI. Scale bar, 25 μm.

## Discussion

Despite many attempts to develop therapeutic drugs for fibrosis, only pirfenidone and nintedanib have been approved for pulmonary fibrosis^15^, but no effective drugs for liver fibrosis have been developed. As HSCs are the major driver of liver fibrosis, HSCs should be an ideal target to develop drugs for liver fibrosis^3,4^. HSCs in the normal liver are quiescent and are activated to produce collagens by various liver insults. Upon removal of liver injury agents, aHSCs undergo cell death or return to a preactivated state^2^. Thus, there are three possible ways to develop anti-fibrotic drugs for liver fibrosis, i.e. blocking activation of qHSCs, induction of cell death of aHSCs, and deactivation of aHSCs^1^. We previously developed a HTS for chemical compounds that suppress the activation of qHSCs derived from hiPSCs, and successfully identified some drug candidates, such as artemisinin as an anti-fibrotic drug^10^. Because drugs to suppress HSCs activation can be used as a prophylactic therapy, it would be more beneficial to develop drugs that reverse aHSCs for a more advanced stages of fibrosis. In this paper, we attempted to reverse aHSCs to daHSCs by using various chemicals that have been reported to play a role for fibrosis. We found that a combination of chemical compounds, SDYLRP, not only suppressed expression of activation marker genes, *ACTA2*, *COL1A1*, but also induced expression of *LHX2* and *LRAT* that are expressed in qHSCs. Importantly, *TCF21*, a transcription factor^9^ that was shown to deactivate aHSCs, was also upregulated by SDYLRP. Moreover, morphological change of HSCs by phalloidin staining also showed reversion of aHSCs to qHSCs by SDYLRP. These results indicate that aHSCs can be reverted to a pre-activation stage by chemicals, encouraging us to set up a HTS.

As HTS requires a large number of aHSCs, we expanded the culture of aHSCs from hiPSCs and cryopreserved the cells. We also proposed a new concept, the Actin score, as a new indicator for screening compounds of aHSC deactivation and established HTS. This SiR-actin staining establishes a simple and versatile screening system to assess deactivation of HSCs without the need for specialized reporter cell lines. Using this system, we screened a chemical library consisting of 3,909 existing chemical drugs and reagents. After 3 rounds of screening, we were able to find 25 compounds that suppressed the expression of *ACTA2* and *COL1A1*, while promoted the expression of *TCF21*, *LRAT* and *LHX2*. Among them, we selected GZD824 that exhibited a strong effect for further study. It is a pan-Bcr-Abl inhibitor that has been used for treatment of chronic myeloid leukemia (CML)^16^. GZD824 dose-dependently suppressed expression of *ACTA2* and *COL1A1* and enhanced expression of *TCF21*, *LRAT*, *MDK* and *PTN*. Furthermore, phalloidin staining showed a dramatic morphological change of aHSCs with actin stress fibers to qHSC-like cell morphology without actin fibers. These results indicate that a single compound can revert, at least in part, the daHSC phenotype to a pre-activation stage, and that our HTS using aHSCs derived from hiPSCs is useful to find an effective drug candidate that can ameliorate an advanced stage of fibrosis.

## Acknowledgements

We thank Hiroko Anzai, Chizuko Koga, Yoshiko Kamiya and Misao Himeno for technical assistance. This study was supported by JSPS KAKENHI (Grant Number 23H03836 and 21H02710), Japan Agency for Medical Research and Development (Grant Number JP21bk0104136 and JP21ek0109502), Basis for Supporting Innovative Drug Discovery and Life Science Research from AMED under Grant Number JP22ama121053 (support number 3355), and ISM Co., Ltd.

## Author contributions

Y.N. designed the study, performed experiments, analyzed data, and edited the manuscript. T.I. and M.T. contributed discussion and interpretation. A.M. and T.K. designed the study and wrote the manuscript.

---

**Figure S1, related to Figure 2.**

(A) Relative gene expression of HSC markers in qHSCs (day 0), aHSCs (day 7 and day 10) and daHSCs. The results are shown as the mean ± SD of independent experiments.

(B) Fluorescence images (Collagen I: green, αSMA: red and phalloidin: white) of qHSCs (day 0) and aHSCs treated with DMSO, RP, LRP, YLRP, SDLRP or SDYLRP. Y: Y27632, L: Lanifibranor, R: Retinoic acid, P: Palmitic acid, S: SB431542, D: Dorsomorphin. Nuclei were stained with DAPI. Scale bar, 50 μm.

**Figure S2, related to Figure 4.**

Dot plots of Actin score per well of 384-well plates (plate number 6036 to 6049) incubated with compounds of the chemical library. Red dots indicate compounds with -3 SD of the mean and green dots indicate compounds with -2 SD of the mean.

**Figure S3, related to Figure 4.**

Relative expression levels of *ACTA2*, *COL1A1*, *TCF21*, *LRAT* and *LHX2* in hiPSC-derived aHSCs incubated with compound A to Y at the concentration of 125 nM, 500 nM and 2 μM. The values of samples treated with DMSO were set to 1. The results are shown as the mean ± SD of independent experiments (each experiment contains three technical replicates).

## Methods

### Induction of qHSCs from hiPSCs

The hiPSC line TkDN4-M provided by Dr. Otsu at the Institute of Medical Science, The University of Tokyo was used in this study and qHSCs were prepared from hiPSCs according to our previous protocol^10^.

### Activation of qHSC-like cells *in vitro* and cryopreservation

hiPSC-derived qHSC-like cells were seeded on Cellmatrix Type I-C (Nitta gelatin)-coated plates at a density of 10,000 cells/cm^2^ in Stempro-34 SFM medium supplemented with Y27632 (10 μM). After 72 h of culture, medium was changed to Stempro-34 SFM medium. HSCs cultured for a total of 7 days were washed with PBS, detached from the bottom of the culture with TrypLE Express/EDTA (Gibco), and collected as single cells. The collected cells were suspended in Cell Banker 1 (Nippon Zenyaku Kogyo) at 4 × 10^6^ cells/mL, dispensed into tubes for cryopreservation at 500 μL/tube, and frozen at -80°C for storage.

### Deactivation induction of aHSCs

Seven days cultured aHSCs were cultured in Stempro-34 SFM medium (Thermo Fisher Scientific) supplemented with SB431542 (5 μM) (Tocris), Dorsomorphin dihydrochloride (0.5 μM) (Tocris), Y27632 (10 μM) (Wako Pure Chemical Industries, Ltd.), Lanifibranor (30 μM), retinol (10 μM) (Sigma), and palmitic acid (10 μM) (Sigma). Cell morphology was evaluated by staining these cells with FITC-conjugated Phalloidin (Molecular Probe) for the cytoskeletal molecule F-Actin.

### Quantitative RT-PCR

Total RNA was isolated from cultured cells using the NucleoSpin RNA XS (MACHEREY-NAGAL, Duren, Germany), according to the manufacture’s protocol. Subsequently, the RNA was reverse transcribed using the PrimeScriptII 1st strand cDNA Synthesis Kit (Takara bio, Shiga, Japan). Quantitative RT-PCR was performed using SYBR Premix EX TaqII (Takara bio) with specific primers listed in Supplementary Table 1. All data were calculated using the ddCt method with *GAPDH* as a normalization control.

### RNA sequence analysis

Total RNA was extracted from qHSCs (day 0), aHSCs (day 7, day 10) and daHSCs (day 10) using the NucleoSpin RNA XS (MACHEREY-NAGAL, Duren, Germany), according to the manufacture’s protocol. Libraries were prepared using the Optimal Dual-mode mRNA Library Prep Kit (BGI). All libraries were sequenced using the DNBSEQ-G400 (MGI).

### Immunofluorescent staining of cultured cells

Cultured cells were fixed with 10% formaldehyde for 20 minutes at 4 °C, then incubated overnight at 4 °C with the specific primary antibodies. They were subsequently incubated with appropriate fluorescent secondary antibodies and examined under a fluorescence microscope (BZ-X810, Keyence). The primary and secondary antibodies used were as follows: Collagen I (Bio-Rad 2150-1410, 1:300), α-SMA (Sigma A2547, 1:500), Alexa Fluor 647 Phalloidin (Thermo fisher A22287, 1:300), Alexa Fluor 488 rabbit IgG (Thermo fisher A21206, 1:400) and Alexa Fluor 555 mouse IgG (Thermo fisher A31570, 1:400).

### Validation of drug screening system

Stempro-34 SFM medium containing 0.2% DMSO was added to collagen-coated 384-well plates at 20 μL/well. Cryopreserved aHSCs were suspended in Stempro-34 SFM medium at 5,000 cells / 20 μL and added to the each well with medium. After 2 days of culture, 20 μL of Stempro-34 SFM medium containing 0.5 μM SiR-Actin (Cytoskeleton, Inc) was added and incubated overnight. F-Actin filaments labeled with Sir-Actin were visualized using BZ-X810 (Keyence) and Actin score was calculated in each well to determine the CV (%) and Z’-factor.

### Drug screening for aHSC deactivation

Validated compound library (3,909 compounds) was provided by Drug Discovery Initiative, The University of Tokyo. Using the same procedure as the validation of drug screening system, Actin score was calculated for each compound at a concentration of 2 μM and 400 nM. The mean and SD of the Actin score for each plate were calculated, the compounds that exhibited Actin score 2 × SD below the mean were selected. To select the lead candidate compounds, quantitative RT-PCR was performed in the hiPSC-derived aHSCs at the concentration of 125 nM, 500 nM and 2 μM.

### Statistics

All experiments were independently repeated using at least three times. Values are expressed as the mean ± SD. Statistical analyses were performed using Microsoft Excel (Microsoft, Seattle, WA) and GraphPad Prism (GraphPad Software Inc., San Diego, CA). Statistical differences between groups were evaluated using the One-Way ANOVA (two-tailed, *vs* DMSO), and *P* values <0.05 were considered statistically significant.

