## Supplementary table for "Development of a high throughput system to screen compounds that revert the activated hepatic stellate cells to a quiescent-like state"

Supplementary Table 1

| | |
|---|---|
| *ACTA2* | CAGCCAAGCACTGTCAGG |
| | CCAGAGCCATTGTCACACAC |
| *COL1A1* | AAGAGGAAGGCCAAGTCGAG |
| | CACACGTCTCGGTCATGGTA |
| *GAPDH* | AAGGTGAAGGTCGGAGTCAA |
| | AATGAAGGGGTCATTGATGG |
| *LHX2* | ATGCTGTTCCACAGTCTGTCG |
| | GCATGGTCGTCTCGGTGTC |
| *LRAT* | TACTGCAGATATGGCACCCC |
| | CCAAGACTGCTGAAGCAAGA |
| *MDK* | CAAGTTTGAGAACTGGGGTGCGTG |
| | AGTCCTTTCCCTTCCCTTTCTTGG |
| *PTN* | CAGCGTCGAAAATTTGCAGCTGC |
| | TCCACTGCCATTCTCCACAGTCAG |
| *TCF21* | CACTTGAGGCAGATCCTGGCTA |
| | CGGTCACCACTTCTTTCAGGTC |
