## Supplementary figures and images for "Development of a high throughput system to screen compounds that revert the activated hepatic stellate cells to a quiescent-like state"

### Supplementary figures

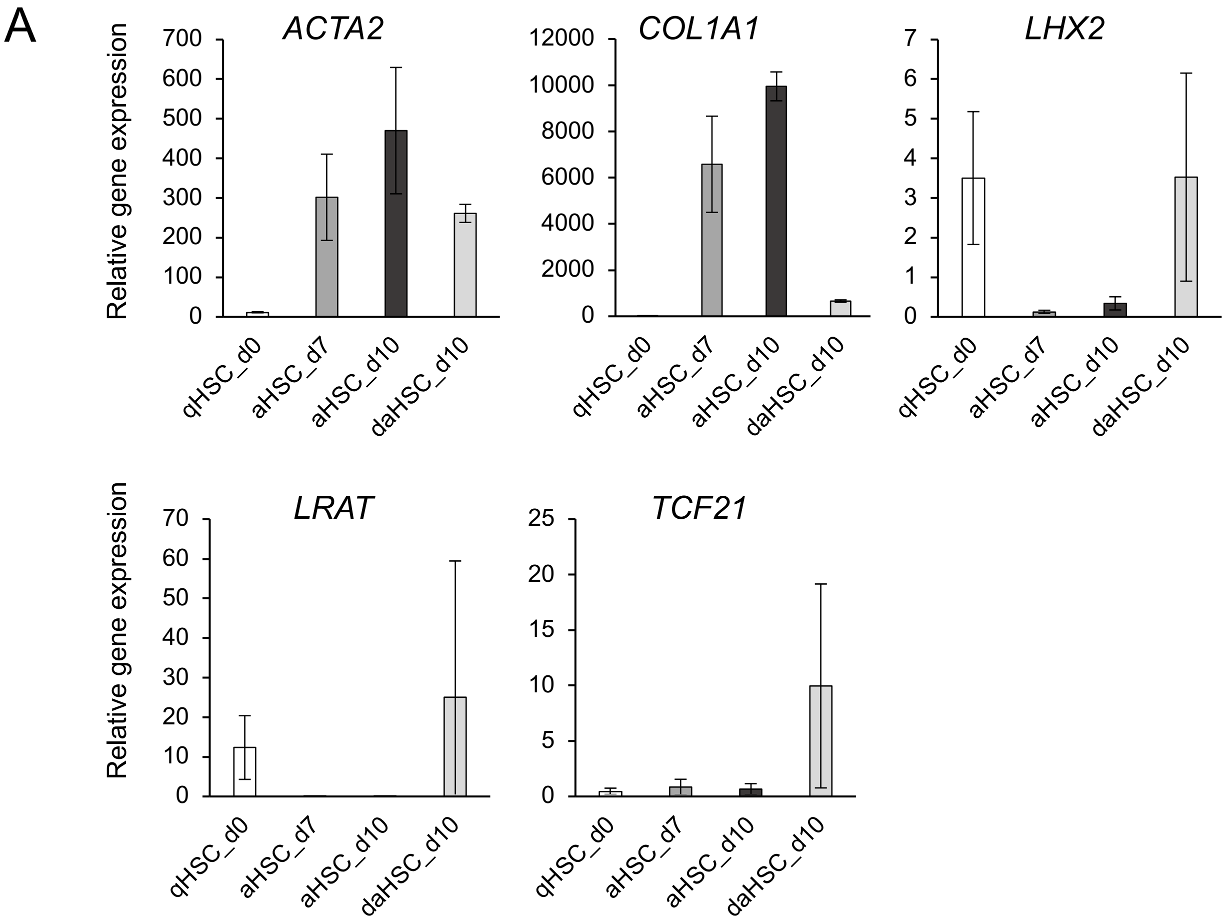

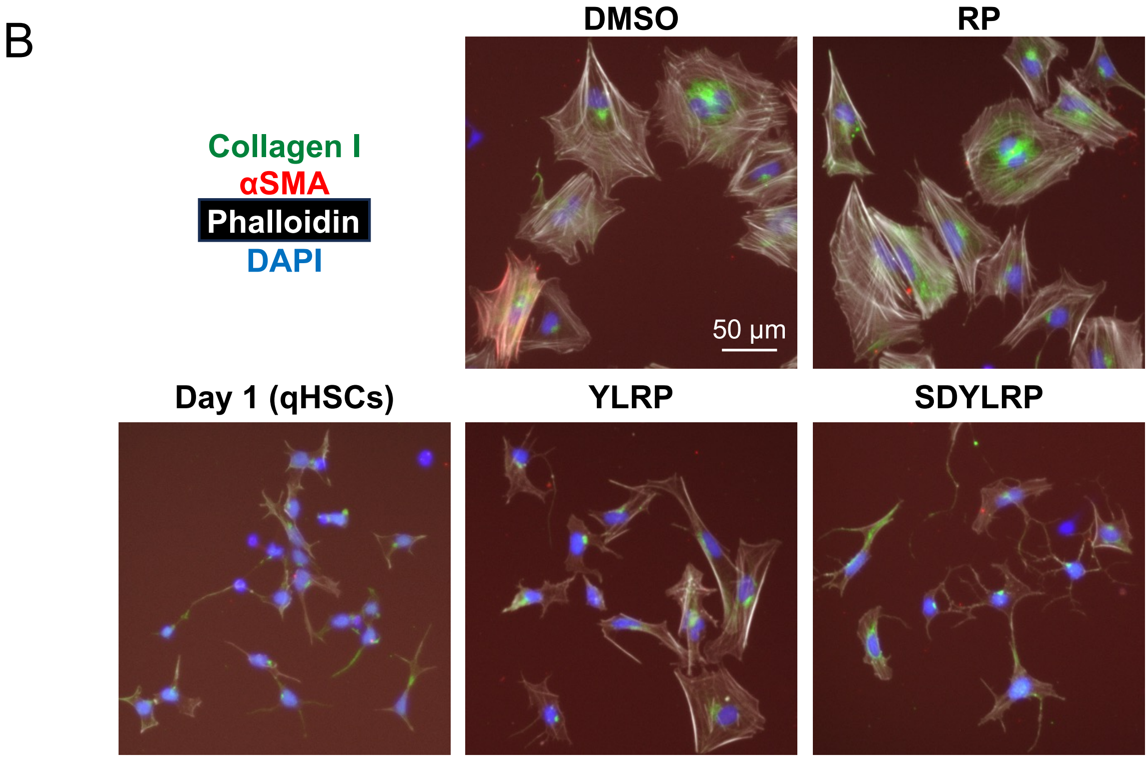

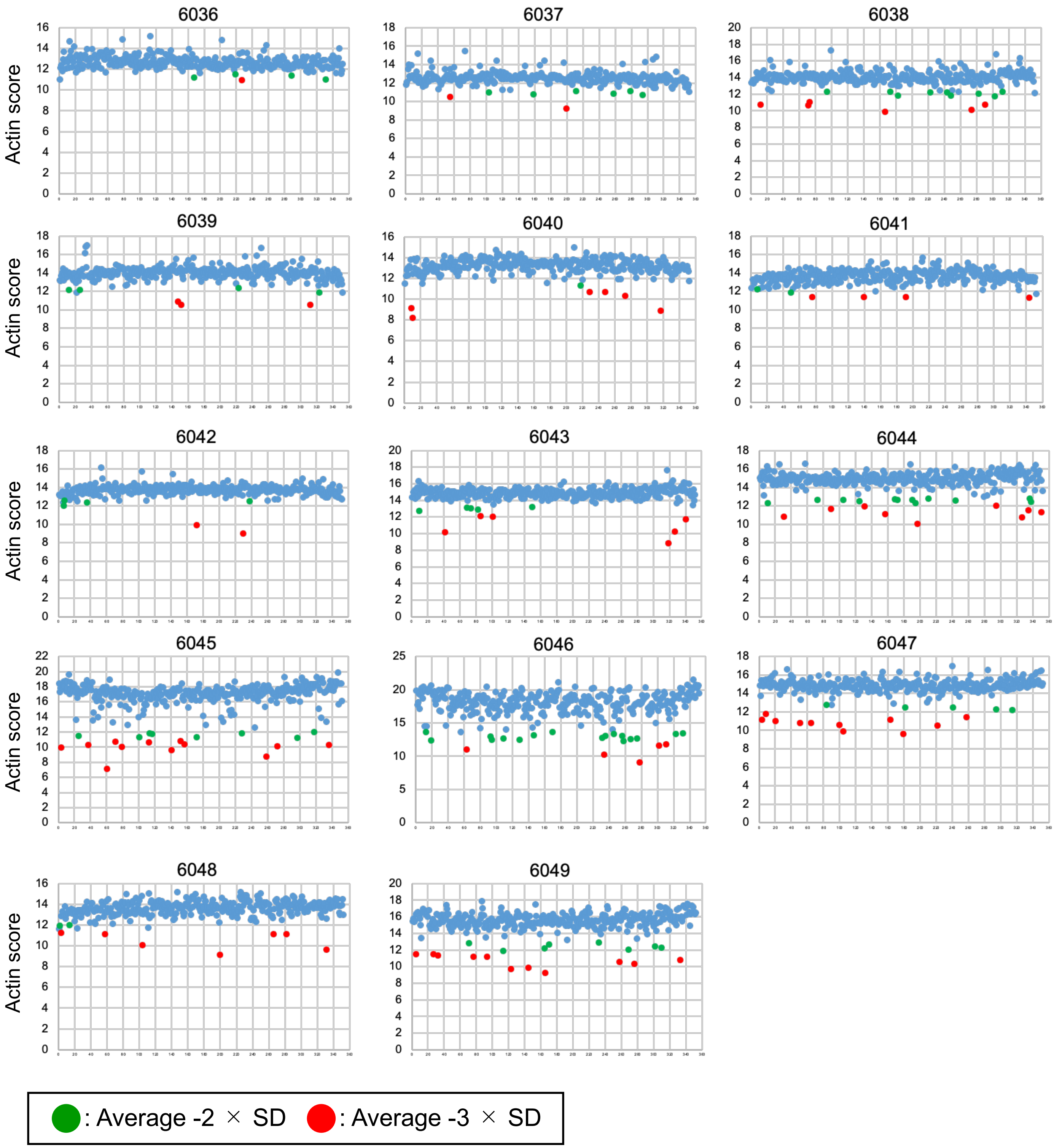

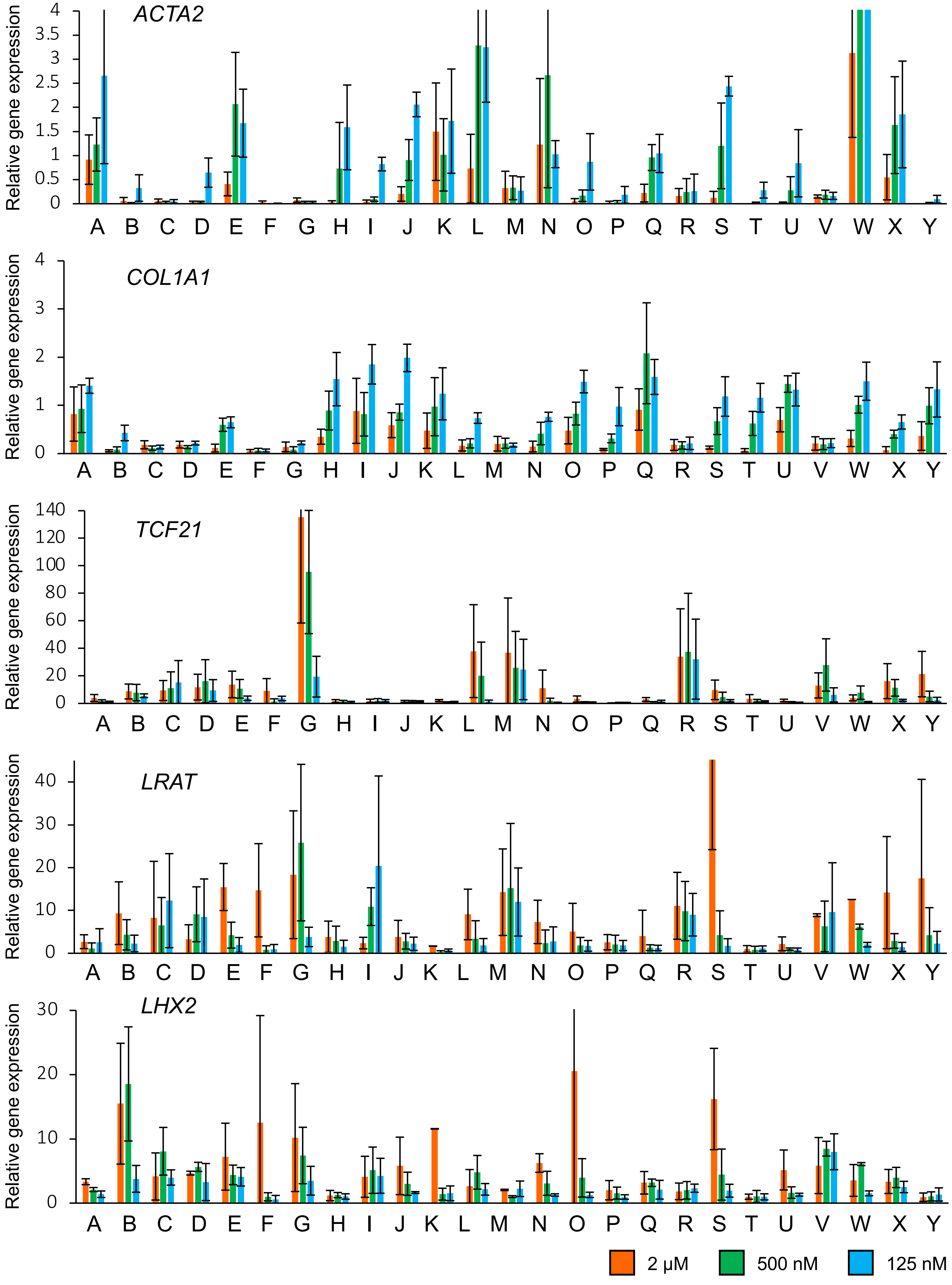
